# Yeast Genomic Screens Identify Kinesins as Potential Targets of the *Pseudomonas syringae* Type III Effector, HopZ1a

**DOI:** 10.1101/365692

**Authors:** Amy Huei-Yi Lee, D. Patrick Bastedo, Timothy Lo, Maggie A. Middleton, Ji-Young Youn, Inga Kireeva, Jee Yeon Lee, Sara Sharifpoor, Anastasia Baryshnikova, Jianfeng Zhang, Pauline W. Wang, Sergio G. Peisajovich, Michael Constanzo, Brenda J. Andrews, Charles M. Boone, Darrell Desveaux, David S. Guttman

**Affiliations:** Department of Cell & Systems Biology, University of Toronto, Toronto, Ontario, Canada; Centre for the Analysis of Genome Evolution & Function, University of Toronto, Toronto, Ontario, Canada; The Donnelly Centre, University of Toronto, Toronto, Ontario, Canada; Lewis-Sigler Institute for Integrative Genomics, Princeton University, Princeton, New Jersey, USA; Department of Molecular Genetics, University of Toronto, Toronto, Ontario, Canada

**Author notes:** To whom correspondence should be addressed (D.S.G.); (D.D.). These authors contributed equally (Amy Huei-Yi Lee, D. Patrick Bastedo). These authors contributed equally (Darrell Desveaux, David S. Guttman).

**Keywords:** *Pseudomonas syringae*, Type III secreted effector, Pathogenic Genetic Array, Yeast screen, Pathogen-host interactions, HopZ1, Kinesin

## Abstract

Gram-negative bacterial pathogens inject type III secreted effectors (T3SEs) directly into host cells to promote pathogen fitness by manipulating host cellular processes. Despite their crucial role in promoting virulence, relatively few T3SEs have well-characterized enzymatic activities or host targets. This is in part due to functional redundancy within pathogen T3SE repertoires as well as promiscuous individual T3SEs that can have multiple host targets. To overcome these challenges, we conducted heterologous genetic screens in yeast, a non-host organism, to identify T3SEs that target conserved eukaryotic processes. We screened 75 T3SEs from the plant pathogen *Pseudomonas syringae* and identified 16 that inhibited yeast growth on rich media and eight that inhibited growth on stress-inducing media, including the acetyltransferase HopZ1a. We focused our further analysis on HopZ1a, which interacts with plant tubulin and alters microtubule networks. We first performed a Pathogenic Genetic Array (PGA) screen of HopZ1a against ~4400 yeast carrying non-essential mutations and found 95 and 10 deletion mutants which reduced or enhanced HopZ1a toxicity, respectively. To uncover putative HopZ1a host targets, we interrogated both the genetic- and physical-interaction profiles of HopZ1a by identifying yeast genes with PGA profiles most similar (i.e. congruent) to that of HopZ1a, performing a functional enrichment analysis of these HopZ1a-congruent genes, and by analyzing previously described HopZ physical interaction datasets. Finally, we demonstrated that HopZ1a can target kinesins by acetylating the plant kinesins HINKEL and MKRP1.

**ARTICLE SUMMARY:** Bacterial pathogens utilize secretion systems to directly deliver effector proteins into host cells, with the ultimate goal of promoting pathogen fitness. Despite the central role that effectors play in infection, the molecular function and host targets of most effectors remain uncharacterized. We used yeast genomics and protein interaction data to identify putative virulence targets of the effector HopZ1a from the plant pathogen *Pseudomonas syringae*. HopZ1a is an acetyltransferase that induces plant microtubule destruction. We showed that HopZ1a acetylated plant kinesin proteins known to regulate microtubule networks. Our study emphasizes the power of yeast functional genomic screens to characterize effector functions.

## INTRODUCTION

Bacterial pathogens of both plants and animals subvert key host processes in order to suppress host immunity and manipulate nutrient supplies. Many Gram-negative bacterial pathogens achieve this goal by delivering type III secreted effectors (T3SEs) into the host cytosol where they manipulate the host in a variety of ways, including modulating signaling pathways, transcription, intracellular transport, cytoskeletal stability, and host defenses (|Büttner and Bonas 2003; Jin et al. 2003; Cornelis 2006; Zhou and Chai 2008; Lewis et al. 2009). Although many bacterial T3SEs have been shown to generally suppress host immunity, we know relatively little about the specific virulence targets and mechanisms of action of most T3SEs. This difficulty in functional characterization of T3SE virulence mechanisms is due to a number of factors, including: (1) redundant targeting of a given host protein by multiple effectors which confounds analysis of individual T3SE deletion mutants; (2) promiscuous individual effectors which can target multiple host proteins, thereby making it difficult to ascribe a virulence function to any individual target (Lewis et al. 2011; Deslandes and Rivas 2012); (3) effectors often show no similarity to proteins or domains with characterized functions, limiting bioinformatic approaches to infer effector functions; and (4) effectors can trigger immune responses as a result of host recognition, which complicates virulence target identification.

In order to gain new insights into the biochemical functions and host targets of bacterial T3SEs, a number of research groups have utilized the model organism *Saccharomyces cerevisiae* (yeast) as a tool (Yoon et al. 2003; Alto et al. 2006; Kramer et al. 2007; Slagowski et al. 2008; Alemán et al. 2009; Munkvold et al. 2009; Salomon and Sessa 2010). The rationale for using yeast to characterize bacterial effectors rests on the fact that many biological processes (for example central metabolism, the control of cytoskeleton dynamics, vesicle trafficking, signal transduction, DNA metabolism and cell cycle processes) are conserved amongst eukaryotes (Siggers and Lesser 2008; Curak et al. 2009; Botstein and Fink 2011; Dolinski and Botstein 2007). Therefore, effectors that target a conserved cellular process in a higher eukaryote may also act on the same cellular process in the simpler and genetically-tractable yeast system. This is particularly attractive if the original host is not readily amenable to high-throughput assays. Another advantage of studying bacterial T3SEs in the yeast system is that the expression of non-effector bacterial proteins does not generally affect yeast growth (Slagowski et al. 2008). This indicates that most fitness defects observed upon T3SE expression in yeast is specifically due to the T3SE, and not simply due to the heterologous overexpression of bacterial proteins. Finally, the expression of translocated effector proteins from both plant and animal pathogens has been shown to inhibit yeast growth by targeting conserved eukaryotic proteins (Munkvold et al. 2008; Siggers and Lesser 2008; Curak et al. 2009; Salomon et al. 2011). For instance, the *Yersinia* T3SE YopJ has been shown to disrupt mammalian innate immunity by preventing the activation of MAPK kinase (MAPKK) and subsequently blocking the MAPK and NFκB signaling pathways (Orth et al. 1999; Orth et al. 2000). Even though yeast cells lack key components of the mammalian innate immune system, YopJ was shown to inhibit MAPK pathways in yeast by preventing the activation of MAPKK as previously observed in mammalian systems (Yoon et al. 2003).

A number of groups have developed yeast genomics tools to characterize bacterial effectors that target conserved eukaryotic cellular processes (Alto et al. 2006; Kramer et al. 2007). A very successful genetic approach is the Pathogenic Genetic Array (PGA), a variation of the well-established Synthetic Genetic Array (SGA) technology, which enables high-throughput genetic screens to identify conserved host targets (Alto et al. 2006; Kramer et al. 2007). The SGA technology involves a series of robotics-assisted cell matings to introduce any marked allele of interest into an array of mutants, allowing the systematic generation of double mutants and the interrogation of di-genic genetic interactions at a genome-wide scale (Tong et al. 2001; Tong et al. 2004; Costanzo et al. 2010). Genetic interactions between two mutations are inferred when the observed double mutant phenotype deviates from the expected phenotype of the combined single mutants. In extreme cases, a synthetic lethal interaction occurs when the combination of two non-lethal mutations causes cell death. Large scale, genome-wide SGA screens have provided genetic interaction profiles for ~75% of the yeast genome (Costanzo et al. 2010). Since genes within the same pathway or bioprocess tend to show very similar genetic interaction profiles, querying the genetic interactions of an unknown gene against the nearly complete SGA compendium of the yeast genome can be a powerful way to predict functions of uncharacterized genes (Costanzo et al. 2010).

Similar to SGA, PGA queries a pathogen effector against a collection of viable yeast deletion strains in a high-throughput array format to analyze effector functions. PGA identifies those yeast deletion mutants that can either reduce or enhance effector-mediated growth defects, and subsequently guides the inference of functional relationships between these yeast genes and the pathogen T3SEs (Alto et al. 2006; Kramer et al. 2007). This PGA strategy was first used to identify yeast deletion mutants that suppress *Shigella* T3SE IpgB2-induced toxicity (Alto et al. 2006). Consistent with the ability of IpgB2 to interfere with Rho1p signaling in mammalian cells, the genetic suppressors of IpgB2 in yeast are downstream of Rho1p, part of the cell wall integrity MAPK-signaling pathway (Alto et al. 2006). Overall this PGA screen revealed that IpgB2 functions as a G protein mimic, capable of activating the Rho1p pathway (Alto et al. 2006).

In this study, we hypothesized that T3SEs that target evolutionarily conserved plant processes can regulate the same processes in yeast. Furthermore, if this conserved process is important for optimal yeast growth, then the overexpression of T3SEs should decrease yeast fitness. We expressed 73 *P. syringae* T3SEs in yeast and identified 24 effectors that reduced yeast fitness, including HopZ1a *_PsyA2_*. In addition, PGA analysis identified yeast genes involved in genetic interactions with HopZ1a; this genetic interaction profile was compared with previously generated SGA datasets to identify yeast genes with interaction profiles similar (or congruent) to that of HopZ1a. In theory, genetic interactors will function in pathways parallel or compensatory to the pathway targeted by HopZ1a. More specifically, yeast genes with genetic interaction profiles congruent to that of HopZ1a may potentially represent direct targets of HopZ1a. Among the yeast genes with HopZ1a-congruent interaction profiles were kinesins, and kinesin homologs from the model plant host *Arabidopsis thaliana* (hereafter *Arabidopsis*) have been previously shown to physically interact with HopZ1a (Mukhtar et al. 2011; Lewis et al. 2012). These findings implicate kinesins as putative targets of HopZ1a and we demonstrated that HopZ1a can acetylate *Arabidopsis* kinesin proteins when co-expressed in yeast. This study emphasizes the power of high-throughput heterologous screens for exploration of T3SE function and for identification of conserved eukaryotic processes that are targeted by diverse pathogens.

## MATERIALS AND METHODS

### Cloning

Promoter-less coding sequences lacking stop codons of *P. syringae* T3SEs were PCR-amplified to include the addition of *att*B1 and *att*B2 linkers and cloned into the Gateway donor vector, pDONR207, using the Gateway BP reactions. T3SEs from *Pto*DC3000, *Psy*B728a and *Pph*1448a were generous gifts from J. Chang (Chang et al. 2005). The additional T3SEs from *Pma*ES4326, as well as T3SEs from the HopZ and HopF families were cloned for this study. The pDONR207-T3SE collection was sequenced-confirmed via Sanger sequencing. These T3SEs were subcloned into the Gateway-compatible yeast integration vector, pBA2262, using the Gateway LR reactions. To confirm the pBA2262-T3SE constructs, purified plasmids were digested with *Bsr*GI or *Not*I and the restriction digest patterns were analyzed.

Promoter-less coding sequences of *A.thaliana* kinesins *HINKEL* (At1g18370) and *MKRP1* (At1g21730) lacking stop codons were likewise PCR-amplified and cloned into pDONR207 and were subcloned by Gateway LR reactions into the autonomously-replicating, single-copy, Gateway-compatible yeast expression vector, pBA350V.

### Yeast strain construction, growth medium, immunoblot analyses

To integrate the P_*GAL1*_-*T3SE-FLAG::NAT^R^* constructs into the yeast genome at the *HO* locus, the SGA query strain (Y7092, MATα, *can1Δ::STE2pr-Sp_his5 lyp1Δ his3Δ1 leu2Δ0 ura3Δ0 met15Δ0*) was transformed with *Not*I-digested BA2262-T3SE plasmid DNA using the standard transformation method (Gietz and Woods 2002).

For immunoblot analyses, yeast strains expressing the FLAG-tagged T3SEs under control of the *GAL1* promoter were grown overnight at 30° shaking (200 RPM) in 1 ml of YP broth with 2% raffinose (YPR) in deep-well plates with sterile glass beads in each well. The overnight cultures were subsequently diluted into deep-well plates containing 1 ml of YP broth with 2% galactose (YPG) at OD600 of 0.1. The cultures were induced for T3SE expression for 7 to 8 hours, or until the cultures reach OD600 of 1. The 1 ml-cultures were pelleted at 13,000 x *g* for 1 min, washed, and frozen at −20°. Whole cell extracts were prepared from TCA-fixed cells as described (Kurat et al. 2009). The protein pellets were resuspended in 1X sample buffer and neutralized by addition of 2M Tris solution. The lysates were separated by 12% SDS-PAGE and immunoblot was performed with mouse anti-FLAG primary antibodies (Sigma, F3165, USA) via chemiluminescence (Amersham, USA).

### Pathogenic genetic array

The pathogenic genetic array (PGA) analysis was based on a variation of the SGA method used for synthetic dosage lethality screens (Tong et al. 2001; Sopko et al. 2006). In brief, Y7092 (the SGA query strain) with integrated *HOΔ::GAL1-hopZ1a-FLAG::NAT^R^* was mated into the 1536-density *MATa* deletion mutant array marked with *KAN^R^*, which represents each single mutant colony four times on the array. Y7092 carrying *HOΔ::NAT^R^* (SN851) was used as a negative control strain. The *MATa/α* diploids were selected on YPD supplemented with clonNAT (100 μg/ml) and G418 (200 μg/ml) at 30° for two days. Diploid cells were pinned onto enriched sporulation media (20 g/L agar, 10 g/L potassium acetate, 1 g/L yeast extract, 0.5 g/L glucose, 0.1 g/L amino acids-supplement) and allowed to sporulate at 22° for at least one week. The spores were pinned onto synthetic dextrose media (SD) – His/Arg/Lys + clonNAT/canavanine/thialysine and incubated at 30° for two days to select for *MATa* haploid meiotic progeny. The drugs canavanine and thialysine were used at 50 μg/ml. The *MATa* haploid meiotic progeny were subsequently pinned onto SD – His/Arg/Lys + clonNAT/ canavanine/ thialysine/ G418 plates twice to select for the final *MATa* meiotic progeny carrying both the *kan^R^* (yeast deletion strains) and *NAT^R^* (*GAL1-T3SE-FLAG* constructs) markers. To induce for T3SE expression, the *MATa* haploid meiotic progeny from final selection were pinned onto the synthetic galactose (SG) media – His/Arg/Lys + clonNAT/canavanine/thialysine/G418, plates were incubated at 30° for two day. We quantified colony sizes using an adapted SGA protocol (Baryshnikova et al. 2010a).

### Confirmation of PGA interactors

Yeast deletion strains that were either putative suppressors or synthetic lethal interactors from the PGA screens were streaked out on YPD with 200 μg/ml of G418 (Invitrogen Life Technologies, USA) and incubated at 30° for 2 – 3 days. Single colonies of each deletion strain were patched onto YPD plates in 1 – 2 cm^2^ patches and incubated at 30° for 1 overnight to allow for actively growing yeast cultures. A single colony of wild type yeast from the deletion array border was also streaked out and patched onto YPD plates as control strains. Each yeast deletion strain was scraped off from the patches (~10^8^ – 10^9^ cells) using sterile toothpicks and arrayed into a 96-well microtiter plate containing 200 μl of sterile water. Yeast cells were washed once with 200 μl of 0.1 M lithium acetate by centrifugation for 5 min at 1,500 x *g* at 20° in a centrifuge with a microtiter plate rotor. Each well of pelleted yeast cells was resuspended with 180 μl of transformation mix (120 μl of 50% w/v PEG-3350, 18 μl of 1 M lithium acetate, and 25 μl of boiled single-stranded carrier DNA). 60 μl each of resuspended cells were subsequently transferred to 96-well microtiter plates containing either 1 μl of purified plasmid DNA pBA350V (empty vector) or 1 μl of purified plasmid DNA (pBA350V*-hopZ1a* and pBA350V*-hopF2*). The remaining 60 μl of cells served as a mock transformation control. The 96-well microtiter plates were incubated at 30° for 30 min followed by heat shock at 42° for 30 min. Cells were harvested by centrifugation for 10 min at 1,500 x *g* at 20° and resuspended in 100 μl of SD. 50 μl of transformed or mock-transformed cells were plated on SD-Leu and incubated at 30° for 3 days. Transformants carrying either pBA350V or pBA350V-T3SE (pBA350V*-hopZ1a* and pBA350V-*hopF2*) were grown on SD-Leu plates and were subsequently used for confirmation by spot dilution assays.

### Spot dilution assay

For spot dilution assay to determine growth inhibition of Y7092 expressing *P. syringae* T3SEs, 1 ml of cultures were grown at 30° and 200 RPM in YPR in deep-well plates that contain sterile glass beads in each well. Ten-fold dilution series of the overnight cultures were spotted onto YPD, YPG, YPD with 1 M sorbitol, YPG with 1 M sorbitol, YPD with 1 M NaCl, or YPG with 1 M NaCl.

For spot dilution assays to confirm the putative PGA hits as either suppressors or synthetic lethal interactors, the deletion strains carrying either the empty vector (pBA350V) or the effector of interest (pBA350V*-hopZ1a)* were grown in synthetic drop-out media lacking Leu with 2% raffinose (SR-Leu) for two overnights at 30° and 200 RPM. The overnight cultures were serially diluted 15-fold and spotted onto SD-Leu, SG-Leu, SD-Leu and 0.5 M NaCl, or SG-Leu and 0.5 M NaCl. Spot dilutions were grown for two to three days before being photographed. Spot assays were quantified using an unbiased visual toxicity score (between 1 to 5), where 1 represented the strongest toxicity (1 spot grew) and 5 represented the least toxicity (all 5 spots grew). A fitness defect score was subsequently calculated using the toxicity score to compare the expected fitness defect to the observed fitness defect of each mutant (Baryshnikova et al. 2010b; Sharifpoor et al. 2012).

### Yeast co-expression, immunoprecipitations and sample preparation

Overnight cultures of yeast strain Y7092 co-expressing FLAG-tagged HopZ1a (wild type or a catalytically-inactive mutant, C216A) with putative acetylation targets MKRP1 or HINKEL were diluted into fresh SD-Leu (2% raffinose) and allowed to grow at 30° for two doublings prior to inducing expression of effector and targets by addition of galactose to a final concentration of 2%. Following 15 h of induction, cultures were mechanically lysed and lysates were incubated with an anti-FLAG agarose resin (Sigma). After washing the resin to remove unbound proteins, FLAG-tagged proteins were eluted by incubating with 100 uL of FLAG peptide solution (150ug/mL FLAG peptide in TBS) for one hour at 4°. Eluted material was dried to a pellet under vacuum and stored at −80° prior to subsequent mass spectrometry analysis. Dried protein samples were re-solubilized in 50 mM ammonium bicarbonate (pH 7.8) and then subjected to reduction with dithiothreitol at 56°, alkylation with iodoacetamide at room temperature, and overnight digestion with sequencing-grade trypsin (Promega, Madison, WI) at 37°. The enzymatic reactions were stopped with 3% formic acid, purified and concentrated with Pierce C18 Spin Columns (Thermo Scientific) and again dried to a pellet under vacuum. Peptide samples were then solubilized in 0.1% formic acid prior to LC-MS/MS analyses.

### LC-MS/MS Analysis of Proteins, Chromatography and Mass Spectrometry

Samples were analyzed on a linear ion trap-Orbitrap hybrid analyzer outfitted with a nano spray source and EASY-nLC 1200 nano-LC system. The instrument method consisted of one MS full scan (400–1400 *m*/*z*) in the Orbitrap mass analyzer, an automatic gain control target of 500,000 with a maximum ion injection of 500 ms, one microscan, and a resolution of 60,000. Six data-dependent MS/MS scans were performed in the linear ion trap using the three most intense ions at 35% normalized collision energy. The MS and MS/MS scans were obtained in parallel fashion. In MS/MS mode automatic gain control targets were 10,000 with a maximum ion injection time of 100 ms. A minimum ion intensity of 1000 was required to trigger an MS/MS spectrum. The dynamic exclusion was applied using an exclusion duration of 145s.

### Protein ID and Database Searching

Proteins were identified by searching all MS/MS spectra against a large *Arabidopsis* database with the addition of the *hopZ* sequences (extracted from the NCBI database) using SEQUEST (Thermo Scientific™ Proteome Discoverer™ software). A fragment ion mass tolerance of 0.6 Da and a parent ion tolerance of 10 ppm were used. Up to two missed tryptic cleavages were allowed. Methionine oxidation (+15.99492 Da), cysteine carbamidomethylation (+57.02146 Da), and acetylation (+42.01057 Da) were set as variable modifications. The generated search results were imported into the Scaffold data analysis platform, an X!Tandem search (Beavis Informatics, Winnipeg, MA) was performed and the peptides were evaluated using a false discovery rate of 0.1% as determined using a reversed version of the database used in the original search. A mzident.xml file was generated from Scaffold and imported into Scaffold PTM (Proteome Software, Portland, OR) to evaluate and score the post translational modifications.

### Data availability

All yeast strains and plasmids described in this study are available upon request. Supplemental Table 1, Supplemental Figures (1-9), and mass spectrometry raw data files are available at Figshare. TableS1.xlsx: congruence scores for yeast genes with genetic interaction profiles similar to that of HopZ1a. FigureS1.tiff: immunoblot analysis of yeast strain Y7092 expressing P. syringae T3SEs. FigureS2.tiff: spot dilution assays to determine growth inhibition profiles of yeast expressing P. syringae T3SEs. FigureS3.tiff: genome-wide phenotypic screens to identify yeast deletion strains that suppressed or were sensitized to P. syringae T3SE expression. FigureS4.pdf: extracted ion chromatograms, reversed phase chromatography and MS/MS spectra supporting identification of two distinct (singly) acetylated forms of the doubly charged HINKEL peptide, VFGPESLTENVYEDGVK. FigureS5.pdf: extracted ion chromatograms, reversed phase chromatography and MS/MS spectra supporting acetylation of the doubly and triply charged MKRP1 peptide, EISCLQEELTQLR. FigureS6.pdf: extracted ion chromatograms, reversed phase chromatography and MS/MS spectra supporting acetylation of the doubly and triply charged MKRP1 peptide, EIYNETALNSQALEIENLK. FigureS7.pdf: extracted ion chromatograms, reversed phase chromatography and MS/MS spectra supporting acetylation of the doubly and triply charged HopZ1a peptide, ELLDDETPSNTQFSASIDGFR. FigureS8.pdf: zoomed-in views of the extracted ion chromatograms presented in Figure S7. FigureS9.pdf: acetylated HINKEL residues are proximal to the kinesin ATP-binding site. FileS1.tar: mass spectrometry raw data files from co-expression of HINKEL and MKRP1 with HopZ1a (HINKEL_HopZ1a_wt.raw, HINKEL_HopZ1a_CA.raw, MKRP1_HopZ1a_wt.raw, MKRP1_HopZ1a_CA.raw).

## RESULTS

### *P. syringae* T3SEs Compromise Yeast Fitness

We performed a fitness-based screen of *P. syringae* T3SEs in yeast to gain insights into the molecular functions of phytopathogenic T3SEs. We screened T3SEs from three widely studied *P. syringae* strains: 22 T3SEs from *P. syringae* pv. *tomato* DC3000 (*Pto*DC3000); 12 T3SEs from *P. syringae* pv. *syringae* B728a (*Psy*B728a); and 17 T3SEs from *P. syringae* pv. *phaseolicola* 1448A (*Pph*1448a). These three strains have finished genome sequences and represent three of the five major *P. syringae* phylogroups (phylogroups 1, 2, and 3, respectively) (Hwang et al. 2005). We also screened 12 T3SEs from *P. syringae* pv. *maculicola* ES4326 (*Pma*ES4326), which belongs to phylogroup 4. Finally, we screened three additional T3SEs from the HopZ family and nine additional T3SEs from the HopF family, as these two effector families are of particular interest to our group (Figure 1) (Ma et al. 2006; Lewis et al. 2008; Wilton et al. 2010).

**Figure 1.**
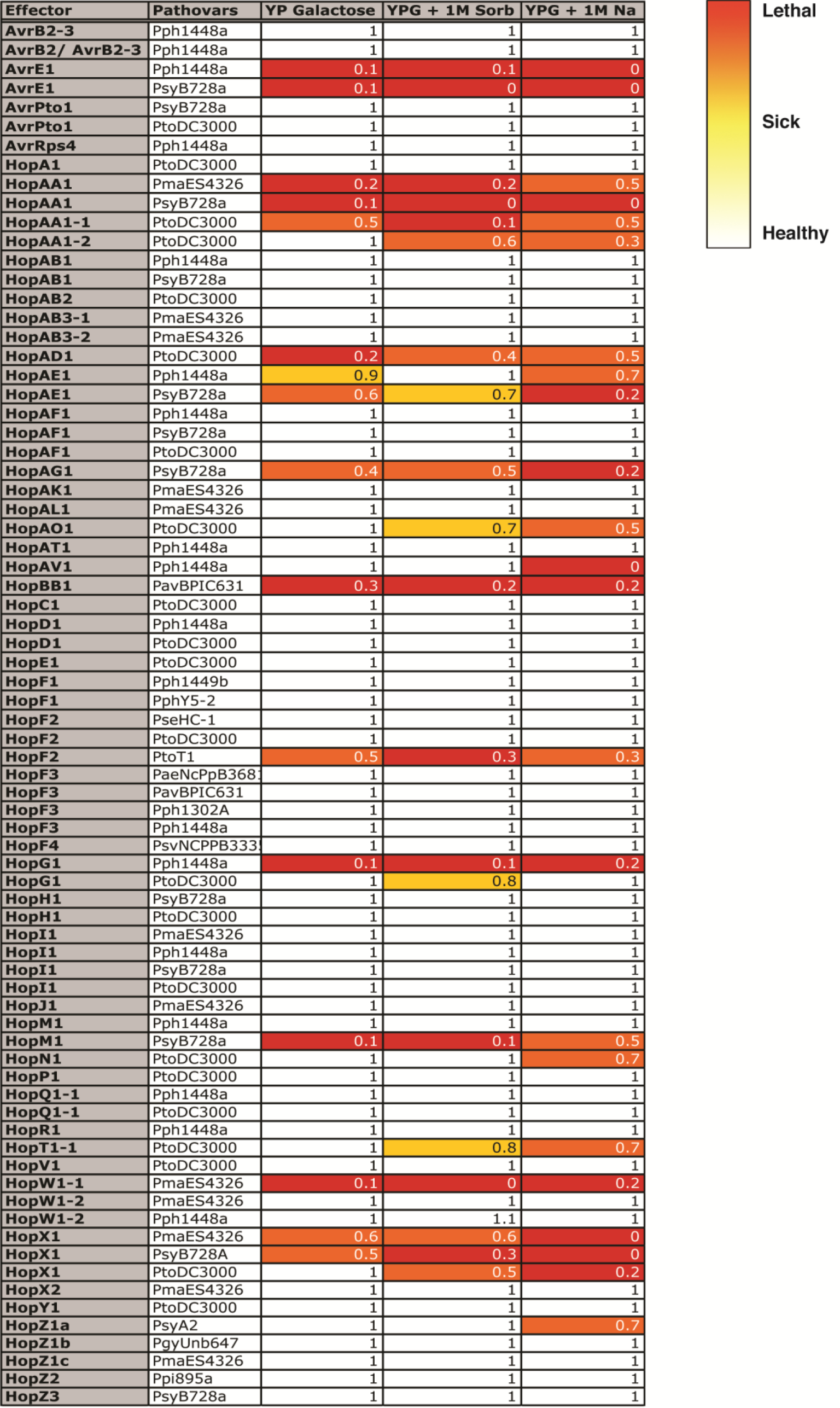
Growth inhibition profiles of yeast (Y7092) expressing 75 *P. syringae* T3SEs on rich media (YP Galactose), rich media with 1 M sorbitol (YPG + 1M Sorb), and rich media with 1 M NaCl (YPG + 1M Na). The growth inhibition by each T3SE at each condition is represented in numbers and heat map, with 1 (or white) corresponding to no growth inhibition to 0 (or red) corresponding to complete growth inhibition. The fitness numbers are calculated for every condition by normalizing the fitness of yeast expressing T3SE to the negative control strain containing the integrated *NAT^R^* antibiotic cassette at the *HO* locus.

For each of these 75 T3SEs, we made yeast strains with chromosomal integrations that introduce a single copy of the T3SE and a drug resistance cassette (*NAT^R^*) at the *HO* locus – a neutral, dispensable locus not functionally required in stable haploid or diploid cells (Baganz et al. 1997). Each T3SE was tagged with a C-terminal FLAG epitope and expressed under the control of a galactose-inducible promoter. We confirmed galactose-dependent expression of 73 T3SEs using western blot analysis (Figure S1).

In order to examine the phenotypic consequence of T3SE expression in yeast, we used serial dilution spot assays on rich media with glucose (T3SE-repressing) and with galactose (T3SE-inducing) media to identify those *P. syringae* effectors that inhibit yeast growth. As expected, we did not observe fitness defects on T3SE-repressing media (Figure S2) compared to the negative control strain (*HO∆::NAT^R^*), however, the expression of 16 out of 73 T3SEs inhibited yeast growth on T3SE-inducing rich media (Figures 1 and S2: AvrE_*Pph1448a*_, AvrE_*PsyB728a*,_ HopAA1*_PmaES4326_*, HopAA1_*PsyB728a*_, HopAA1-1_*PtoDC3000*_, HopAD1*_PtoDC3000_*, HopAE1*_Pph1448a_*, HopAE1*_PsyB728a_*, HopAG1*_PsyB728a_*, HopBB1*_PavBPIC631_*, HopF2*_PtoT1_*, HopG1*_Pph1448a_*, HopM1*_PsyB728a_*, HopW1-1*_PmaES4326_*, HopX1*_PsyB728a_* and HopX1*_PmaES4326_*).

To identify additional T3SEs that may target conserved cellular processes under stress conditions, we also performed fitness assays on media inducing hyperosmotic stress (containing 1 M sorbitol or 1 M NaCl). Eight additional T3SEs altered yeast fitness when induced in the presence of high osmolytes (Figures 1, S2 and S3). Four of the *Pto*DC3000 T3SEs caused enhanced fitness defects in yeast both with 1 M sorbitol and with 1 M NaCl (HopAA1-2, HopAO1, HopT1-1, and HopX1). Although 1 M sorbitol and 1 M NaCl both activate the high osmolarity glycerol (HOG) pathway by creating a high osmolarity environment, NaCl stress creates additional toxicity by altering the ion homeostasis in the cell (Giaever et al. 2002). We also identified a single T3SE that affected yeast fitness only in the presence of 1 M sorbitol (HopG1*_PtoDC3000_*) and three T3SEs that altered yeast fitness only in the presence of 1 M NaCl (HopAV1 *_Pph1448a_*, HopN1*_PtoDC3000_* and HopZ1a*_PsyA2_*).

### Identifying genetic interactors of HopZ1a by PGA analysis

To further characterize *P. syringae* T3SE functions and their mechanisms of toxicity in yeast, we utilized the yeast PGA functional genomics approach. We applied a PGA screen to the T3SE HopZ1a*_PsyA2_* (hereafter HopZ1a), which we have previously shown can bind to tubulin and can alter microtubule networks *in planta* (Lee et al. 2012). We therefore sought to further investigate the role of HopZ1a in regulating microtubule dynamics (and potentially other processes as well) by examining its genetic interaction network in yeast.

To this end, we performed a PGA screen by crossing our integrated HopZ1a-expressing strain with ~4400 haploid yeast non-essential gene deletion mutants (Giaever et al. 2002), looking for those null alleles that either reduced or enhanced HopZ1a-toxicity (Figure S3C). By comparing the colony size of each deletion mutant on glucose (T3SE-repressing) or galactose (T3SE-inducing) media, we classified those deletion mutants with increased or decreased colony size compared to a control array as either suppressors or negative genetic interactors of HopZ1a activity, respectively. The control array was constructed by crossing the same array of ~4400 deletion mutants with a query strain carrying only the *NAT^R^* drug resistance cassette at the *HO* locus (no integrated T3SE). This allowed identification of yeast deletion mutants that were sensitive to galactose and/or NaCl even in the absence of HopZ1a expression. In addition, each array plate of haploid deletion strains contained a border of wild-type yeast carrying the necessary selectable markers for the experimental procedure to correct for border effects. Lastly, to ensure that the expression of HopZ1a did not inhibit yeast mating or sporulation, all of the strain construction steps utilized glucose-containing media to repress HopZ1a expression.

Given that wild type HopZ1a only inhibited yeast growth at high osmolarity (1 M NaCl), we assessed the fitness of ~4400 deletion mutants carrying *GAL-hopZ1a* on media containing galactose and various concentrations of NaCl. At 0.25 M and 0.5 M NaCl, we initially identified 137 deletion mutants with reduced HopZ1a toxicity (suppressors) and 53 deletion mutants with enhanced HopZ1a toxicity (negative genetic interactors; data not shown). To confirm the phenotypes of these 190 deletion mutants, we conducted a secondary screen by transforming the same haploid yeast deletion strains with a single copy plasmid (p*BA350V)* carrying *GAL-hopZ1a*, and then used spot dilution assays to characterize fitness. After excluding those strains with deletions in dubious open reading frames (Winzeler et al. 1999; Giaever et al. 2002; Kramer et al. 2007) and galactose metabolism genes, a remaining 95 suppressors and 10 negative genetic interactors were confirmed by independent transformation and spot dilution assays on 0.5 M NaCl and galactose (Figure 2).

**Figure 2.**
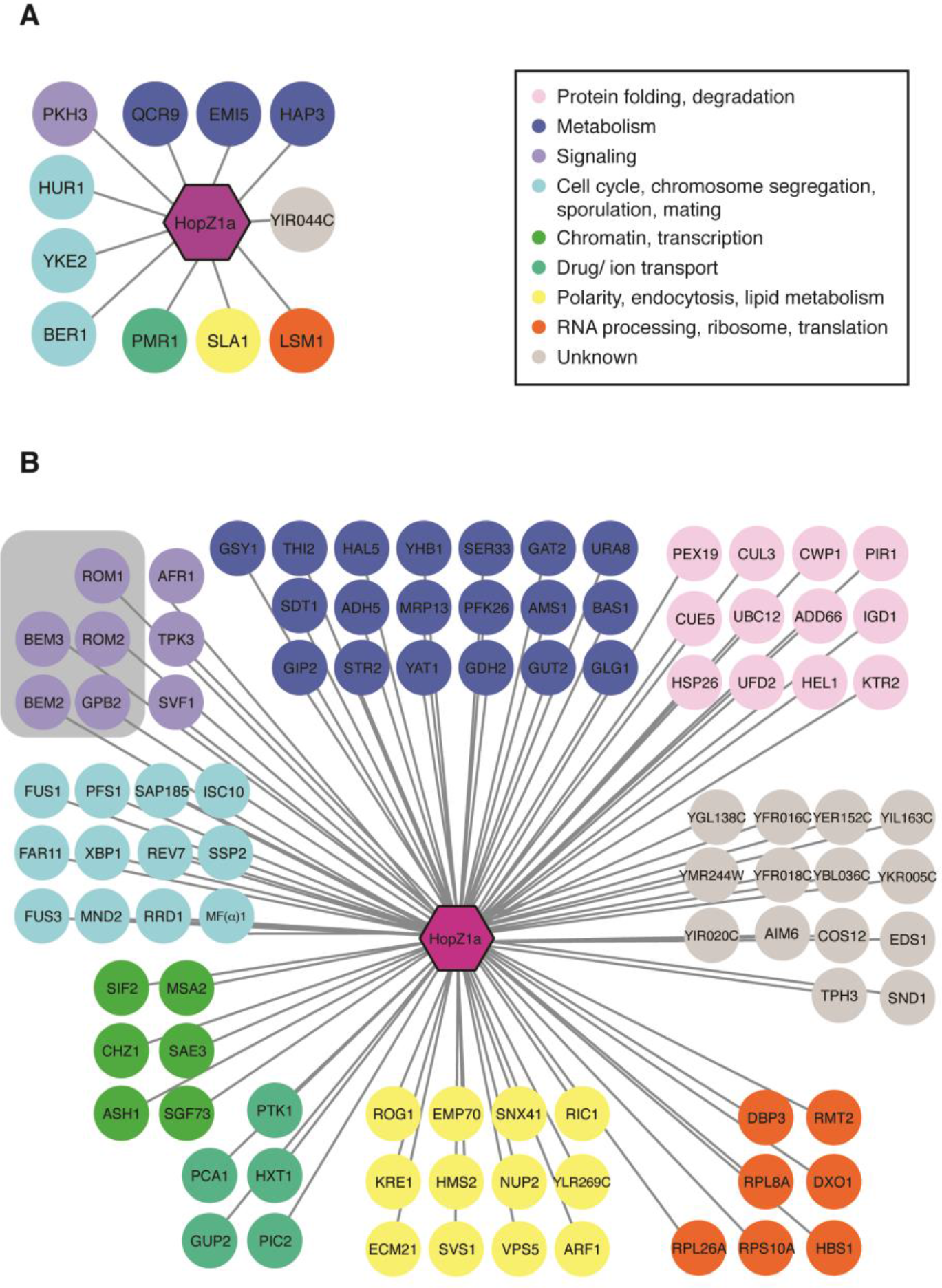
Analysis of HopZ1a suppressors and negative genetic interactors in yeast. Diagram showing (**A**) negative genetic interactors, and (**B**) suppressors of HopZ1a generated using Cytoscape. Nodes are colour coded based on annotations of biological processes from Constanzo *et al*. (Costanzo et al. 2010). The HopZ1a suppressors showed an enrichment in GTPase-mediated signal transduction (gray-shaded box; *p*=0.006).

### Biological processes enriched in HopZ1a PGA profiles

We utilized the Gene Ontology (GO) vocabulary to identify biological processes associated with the confirmed HopZ1a PGA interaction partners, since GO processes that are enriched within this genetic interaction data set may potentially illuminate functional processes that are influenced by HopZ1a (Kramer et al. 2007; Baryshnikova et al. 2010b). We analyzed HopZ1a suppressors using the *Saccharomyces* Genome Database (SGD) GO Term Finder (Hong et al. 2008) and found a significant enrichment of genes involved in GTPase-mediated signal transduction and its regulation (Figure 2B; p=0.006). Specifically, we identified two Rho GTPase activating proteins that are critical for cell polarity and cell division: BEM2 and BEM3; as well as two GDP/GTP exchange proteins: ROM1 and ROM2.

We did not identify significant enrichment of GO processes in the HopZ1a negative genetic interactors. However, two negative genetic interactors of HopZ1a, YKE2 and BER1, are involved in regulating tubulin folding and microtubule-related processes (Figure 2A). Additionally, we identified both suppressors (BEM2, BEM3 and RRD1) and negative genetic interactors (SLA1) that are involved in regulating the actin cytoskeleton (Figure 2A and B). Actin and microtubule cytoskeletons are both involved in fundamental processes such as cell division and intracellular trafficking, raising the intriguing possibility that our genetic interaction screen identified genes whose functions influence both of these two important cytoskeletal components.

### Predicting HopZ1a targets by congruence analysis of genetic interactors

Analysis of the HopZ1a genetic interactors revealed several biological processes that may be disrupted by HopZ1a, yet this information provided limited insight regarding its direct targets. We therefore sought to predict direct targets by identifying yeast gene disruptions that show similar (i.e. congruent) genetic interaction profiles to HopZ1a. This approach is similar to one used previously to identify drug targets in yeast (Costanzo et al. 2010) and assumes that if HopZ1a activity disrupts a given target protein’s function in yeast, the HopZ1a PGA profile would be similar (or ‘congruent’) to the SGA profile of the corresponding gene knockout strain lacking this putative target (Figure 3A and B). We focused our congruency analyses on negative genetic interactors since previous work has indicated that these interactions are easier to interpret than suppressors (Ye et al. 2005).

**Figure 3.**
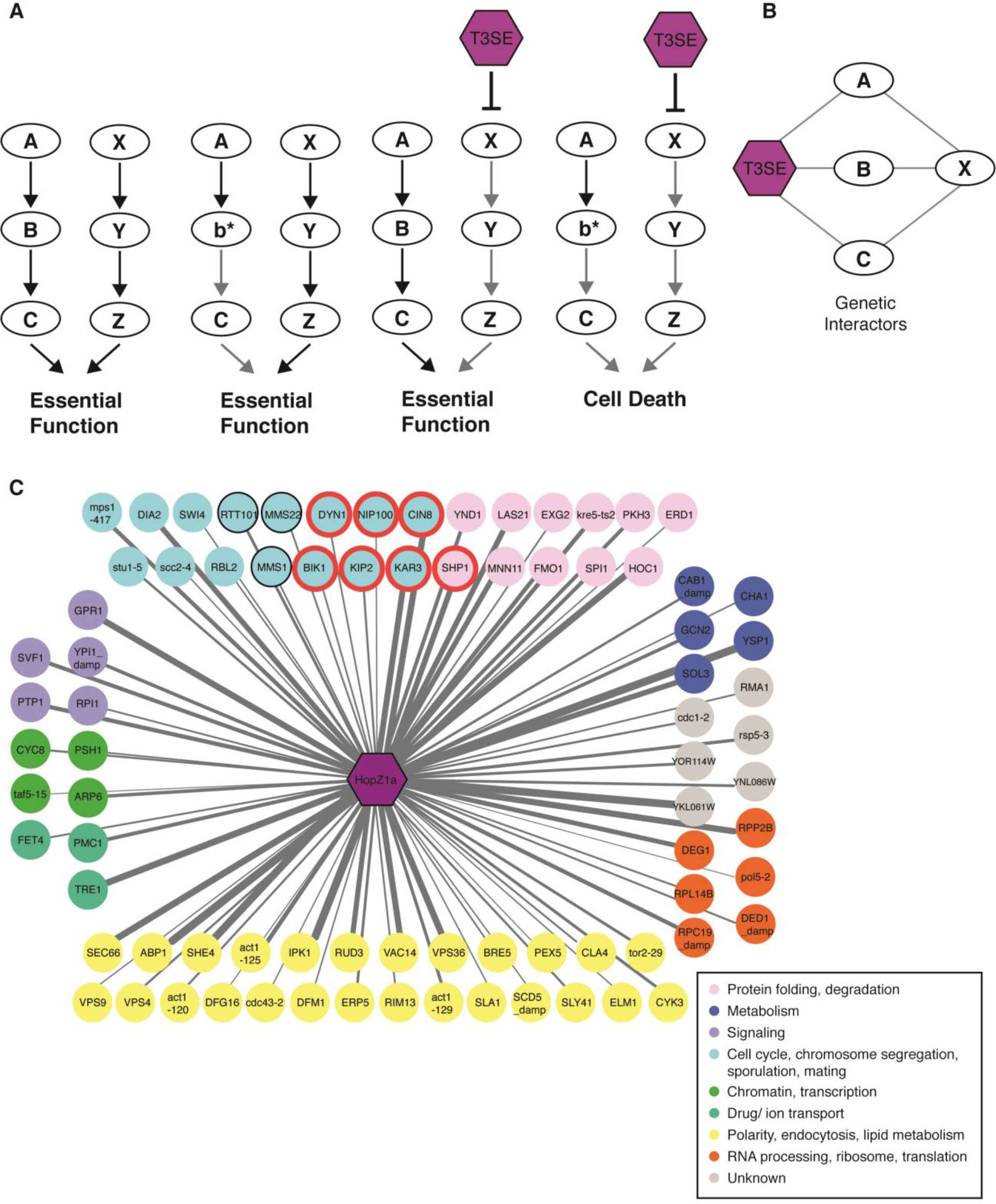
Congruence gene analyses of HopZ1a negative genetic interactors identify microtubule motor proteins as potential targets. (**A**) A model for the molecular mechanism of enhancing T3SE toxicity by targeting redundant pathways. A mutation in either one of the parallel redundant pathways (b* or the inhibition of X by T3SE) does not alter cell viability. However, when both pathways are disrupted (b* and the inhibition of X by T3SE), the cells are not viable. (**B**) Congruence analysis predicts potential T3SE targets by identifying yeast genes (gene X) with similar genetic interaction profiles as the T3SE. (**C**) 81 congruent yeast genes with congruence score ≥ 2 are shown, with nodes color coded based on annotations of biological processes from Constanzo *et al*. (Costanzo et al. 2010). HopZ1a is congruent to yeast deletion strains that are enriched for replication fork processing (*p* < 0.0001) and microtubule-based processes (with a *p* < 0.0004) as analyzed by GOrilla tool (Eden et al. 2009). Genes enriched in microtubule-based processes are circled in red, and genes enriched for replication fork processing are circled in black. Edge thickness is proportional to congruence scores.

To identify yeast genes with HopZ1a-congruent genetic interaction profiles, we compared our HopZ1a genetic interaction profile with those of 1712 single yeast mutants (encompassing ~170,000 interactions) and calculated the pairwise overlap of genetic interactions (Costanzo et al. 2010) using a previously established congruence score (Ye et al. 2005). For any particular congruent gene pair, the significance of the overlap increases with increasing congruence score (Table S1). We identified 99 yeast genes with congruence scores ≥ 2, indicating similarity to the negative genetic interaction profile of HopZ1a (Figure 3C and Table S1). This set of yeast genes with congruent interaction profiles was significantly enriched for genes involved in replication fork processing (p<0.0001) and for genes involved in microtubule-based processes (p<0.0004) (Figure 3C). We were particularly interested in this second group of genes, which includes several microtubule-directed motor proteins such as kinesins (i.e. CIN8, KIP2, VIK1, and KAR3) and proteins in the dynein-dynactin complex (i.e. NIP100 and DYN1) (Figure 3C).

### Bridging genetic and physical interaction data

Given that the natural arena for HopZ1a activity is within plant cells, we examined our set of yeast genes showing genetic interactions with HopZ1a for functional overlap with datasets from two previous studies that had identified direct physical interactions between *A. thaliana* genes and *P. syringae* T3SEs by using yeast two-hybrid screens (Mukhtar et al. 2011; Lewis et al. 2012). Of note, *Arabidopsis* kinesins were identified as HopZ-interacting proteins in both of these previous studies. This overlap between the genetic and physical interactions observed for HopZ1a motivated further investigation into whether *Arabidopsis* kinesins represent direct targets of HopZ1a activity.

### HopZ1a acetylates plant kinesins

Kinesins are microtubule-based motor proteins involved in many cellular processes, including intracellular transport, mitotic cell division, signaling, and microtubule organization (Zhu and Dixit 2012). The kinesins previously shown to participate in direct physical interactions with HopZ family members are the mitochondrially-localized MKRP1 (At1g21730) and MKRP2 (At4g39050) (Mukhtar et al. 2011; Lewis et al. 2012). Since mitochondrial localization of HopZ1a has not been observed (Lewis et al. 2008), we also investigated whether HopZ1a may target related kinesins belonging to the same family as MKRP1 and 2 (i.e. the Kinesin 7 family) (Zhu and Dixit 2012). Of particular interest was the *A. thaliana* HINKEL protein (also known as AtNACK1 or HIK), which is involved in regulating microtubule stability in plants (Strompen et al. 2002; Takahashi et al. 2010; Komis et al. 2011).

HopZ1a is an acetyltransferase with multiple eukaryotic targets, including tubulin and the *A. thaliana* pseudokinase ZED1 (Lee et al. 2012; Lewis et al. 2013). To test whether HopZ1a acetylates kinesins *in vivo*, we used liquid chromatography tandem mass spectrometry (LC-MS/MS) to identify acetylated peptides of both HINKEL and MKRP1. As previously described for the *Arabidopsis* pseudokinase, ZED1 (Lewis et al. 2013), we co-expressed HopZ1a in yeast with each candidate kinesin, all as FLAG-tagged recombinant proteins. LC-MS/MS analysis of anti-FLAG immunoprecipitates identified acetylated peptides from both kinesins (mass increases in multiples of 42 Daltons) present when co-expressed with wild type HopZ1a but not with the catalytically inactive mutant, HopZ1a^C216A^ (Figure 4). Candidate acetylation sites were confirmed by manual inspection of extracted ion chromatograms, reversed phase chromatography and MS/MS spectra (Figures S4-S8 and data not shown).

**Figure 4.**
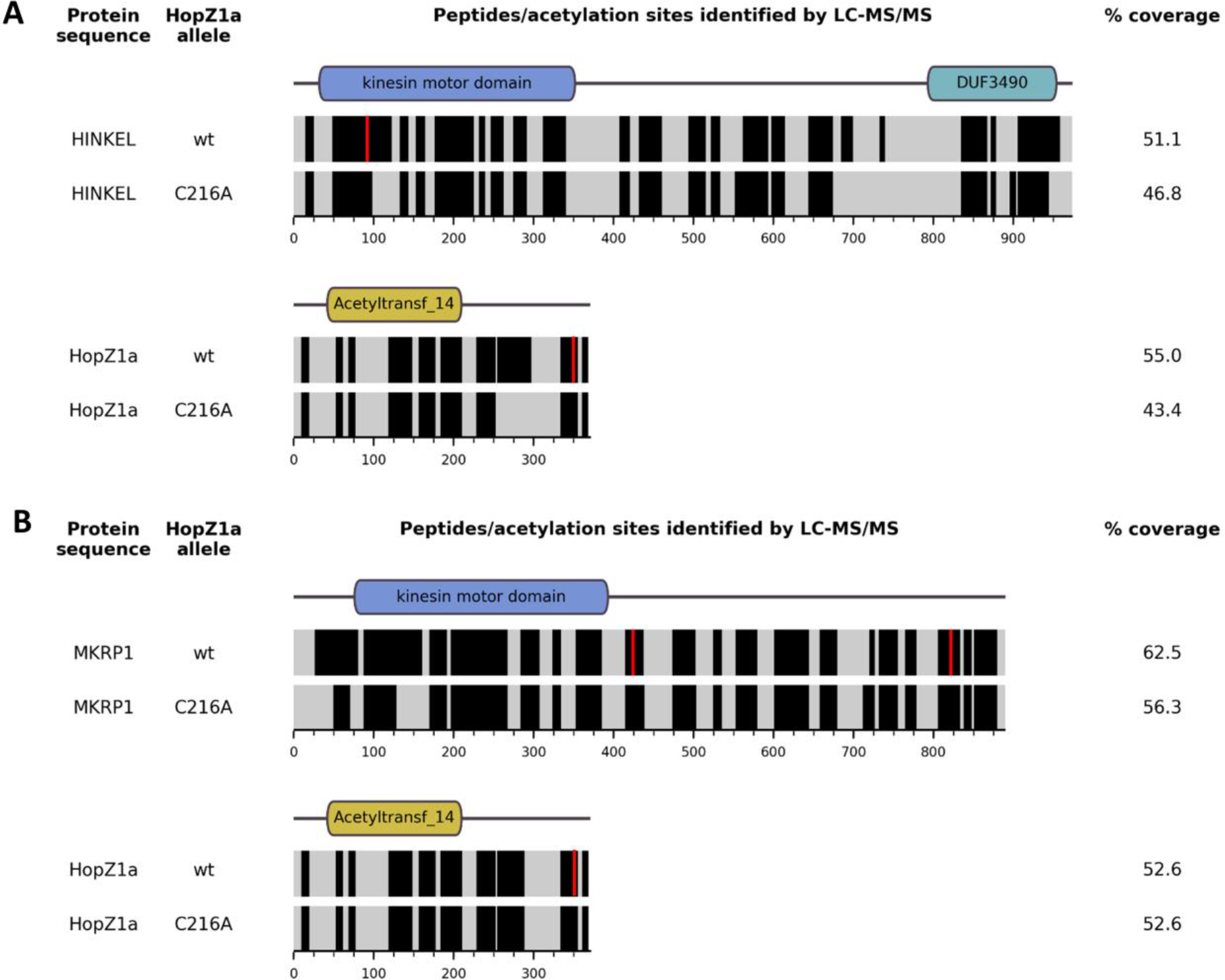
HopZ1a acetylates A. thaliana kinesins HINKEL and MKRP1 when co-expressed in yeast. Predicted domain architectures (as annotated by the NCBI Conserved Domain Database; (Marchler-Bauer et al. 2015) for HINKEL (A), MKRP1 (B) and HopZ1a (A and B) are indicated above horizontal bands representing the mass spectrometry sequence coverage for each protein. Black bands indicate sequences identified with high confidence, while grey bands indicate sequences that were not reliably detected. Red vertical stripes indicate the position of acetylated residues.

In this way we identified two distinct acetylated species of the same HINKEL peptide (VFGPESLTENVYEDGVK; residues 83-99) - ‘peptide A’ (VFGPE[S-Ac]LTENVYEDGVK, acetylated at S88) and ‘peptide B’ (VFGPESL[T-Ac]ENVYEDGVK, acetylated at T90) (Figures 4A, S4). We also identified acetylated peptides from two distinct sites in MKRP1 - ‘peptide C’ (EISCLQEEL[T-Ac]QLR; residues 416-428; acetylated at T425) and ‘peptide D’ (EIYNE[T-Ac]ALNSQALEIENLK; residues 815-33; acetylated at T820) (Figures 4B, S5 and S6). Similar analysis of the HopZ1a-derived peptides from those cells co-expressing MKRP1 indicates auto-acetylation of HopZ1a at three sites in close proximity (T342, S344, T346) (Figures 4, S7, S8) and we did not observe acetylation of HopZ1a in samples from yeast expressing HopZ1a^C216A^. As for cell co-expressing HINKEL and HopZ1a, we identified HopZ1a peptides that represent distinct modifications of the same amino acid sequence (ELLDDETPSNTQFSASIDGFR; residues 336-356) – ‘peptide E’ was singly acetylated at S344 (ELLDDETP[S-Ac]NTQFSASIDGFR), while ‘peptide F’ was doubly acetylated at T342 and T346 (ELLDDE[T-Ac]PSN[T-Ac]QFSASIDGFR). These findings are consistent with a recent report that also described auto-acetylation of T346 (Ma et al. 2015).

## DISCUSSION

In this study, we took advantage of the genetic tractability of yeast to identify putative targets of the T3SE HopZ1a from the plant pathogen *P. syringae*. By integrating and expressing 73 *P. syringae* T3SEs in yeast, we identified 24 effectors that altered yeast fitness on rich media or under high osmolarity conditions, including HopZ1a. We then used a high-throughput PGA screen and analysis of physical interaction datasets to identify kinesin targets of the T3SE, HopZ1a.

Previous studies have identified bacterial phytopathogen T3SEs that altered yeast fitness (Munkvold et al. 2008; Salomon et al. 2011). Of the 27 *Pto*DC3000 effectors tested by Munkvold *et al*., 7 inhibited yeast growth (Munkvold et al. 2008; Munkvold et al. 2009). We tested 20 of these same 27 *Pto*DC3000 effectors and observed fitness phenotypes consistent with these previous data in all cases except for HopAO1, HopD1 and HopN1 (Munkvold et al. 2008; Munkvold et al. 2009). While we integrated T3SEs into the yeast genome and expressed them as single copy genes, Munkvold *et al*. expressed T3SEs on a high-copy plasmid. Differences in gene dosage may thus be contributing to the three instances where our data diverge from this previous report.

Our initial screen provides numerous interesting leads for further study. Notably, *P. syringae* T3SEs encoded in the conserved effector locus (CEL) caused severe fitness defects in yeast (Figure 1). T3SEs of the CEL are conserved across most *P. syringae* strains and typically include the evolutionarily unrelated T3SEs AvrE, HopM1, and HopAA1 (Alfano et al. 2000). *Pph*1448a has nonfunctional alleles of HopM1 and HopAA1 (Joardar et al. 2005), while *Pto*DC3000 contains an additional effector in its CEL, HopN1 (O’Brien et al. 2011). The CEL has been shown to play an important role in bacterial virulence (Alfano et al. 2000; Badel et al. 2003; Munkvold et al. 2009) and in the suppression of salicylic acid (SA)-mediated basal immunity (DebRoy et al. 2004). However, with the exception of HopM1 (Nomura et al. 2006; Nomura et al. 2011), the host targets and the mechanisms by which T3SEs in the CEL promote virulence are not well characterized. Our results suggest that most CEL T3SEs may have evolved to target conserved components of eukaryotic processes. The CEL T3SE-induced yeast fitness defects observed in this study will provide an important tool to help identify virulence targets of this ubiquitous class of phytopathogen T3SEs.

The PGA approach can be used to infer the function of T3SEs by identifying those yeast genes whose deletions either suppress or enhance T3SE lethality. Intuitively, deletion strains that suppress T3SE lethality (known as suppressors) can reveal genes involved in the same pathways as putative T3SE targets. This can be particularly informative when the T3SE activates a pathway resulting in toxicity, as was observed with the Shigella T3SE IpgB2 which activates the Rho1p GTPase signaling pathway in yeast (Alto et al. 2006). However, one caveat of the suppressor screen is that we may identify mutants that suppress T3SE lethality by a general mechanism (i.e. by induction of a general stress response); such genes are unlikely to be informative for the inference of T3SE function.

Deletion mutants that exacerbate the fitness cost of T3SE activity can be explained by either of two alternate mechanisms resulting in ‘negative genetic interactions’. In one case, the T3SE acts in the same pathway as the ‘negative genetic interactor’, resulting in cumulative insults to an essential pathway or complex (Boone et al. 2007; Dixon et al. 2009). Alternatively, the T3SE and ‘negative genetic interactor’ may act on parallel pathways, which redundantly contribute to an essential function (Figure 3A) (Boone et al. 2007; Dixon et al. 2009). Our analysis of both suppressors and negative genetic interactors revealed enrichment of signal transduction pathways involving small-GTPases and may reflect an ability of HopZ1a to influence these cellular processes. However, in order to gain further insight into the direct targets of HopZ1a we also applied congruence analysis to compare SGA interaction profiles of 1712 yeast deletion mutants (Costanzo et al. 2010) with the HopZ1a PGA interaction profile described in this study. This approach is conceptually similar to the integration of chemical-genetic and SGA datasets for identification of drug targets (Costanzo et al. 2010); functional inhibition of a target protein by drug or by T3SE is expected to mimic the consequences of the corresponding target gene’s deletion, resulting in similar/congruent genetic interaction profiles.

Applying these principles, we identified SGA profiles that were most similar to the HopZ1a PGA profile and analyzed them for functional enrichment. Genes involved in replication fork processing (*p* < 0.0001) and microtubule-based processes (*p* < 0.0004) were enriched in the subset with HopZ1a-congruent interaction profiles. We were particularly interested in microtubule-associated processes since HopZ1a can disrupt microtubules in plants and interacts with tubulin in both plant and animal cells (Lee et al. 2012). Indeed, kinesins (known microtubule-guided motor proteins) were identified not only through our congruency analysis, but also by two independent yeast-two hybrid screens for *Arabidopsis* proteins that bind to related HopZ family members. The fact that kinesins are found at the intersection of these three independent datasets indicates that members of this family may indeed represent *bona fide*, direct targets of HopZ1a. In support of this possibility, HopZ1a can acetylate both of the *Arabidopsis* kinesins HINKEL and MKRP1 (Figure 4).

The acetylated sites (S88, T90) of HINKEL are found within its kinesin motor domain (Figure 4A), and mapping these to the corresponding positions in the structure of human kinesin CENP-E (Garcia-Saez et al. 2004) reveals a close proximity to the nucleotide-binding pocket (Figure S9). In *A. thaliana*, HINKEL activates the ANP1/ANQ1/MPK4 MAPK pathway that ultimately regulates microtubule-bundling proteins (e.g. MAP65) via phosphorylation (Komis et al. 2011). Our data suggest a possible mechanism for HopZ1a-mediated antagonism of this pathway whereby nucleotide binding and/or hydrolysis activity is altered following acetylation of sites proximal to the nucleotide-binding pocket of HINKEL.

Although HopZ1a has not been detected in mitochondria, we cannot rule out the possibility that the mitochondrial kinesins identified by yeast two-hybrid assays are also targeted by HopZ1a, especially considering that they are targeted by the *P. syringae* T3SE HopG1 and are involved in plant immunity (Shimono et al. 2016). HopZ1a acetylates MKRP1 at two distinct sites: T425 is just ‘downstream’ of the kinesin motor domain while T820 is near its C-terminus (Figure 4B). In *Nicotiana*, the HINKEL ortholog NACK1 is phosphorylated near the C-terminus at residues T675, T690 and T836 by cyclin-dependent kinases to regulate microtubule dynamics during cytokinesis (Sasabe et al. 2011). Although reasonable speculation might suggest that C-terminal acetylation could disrupt hypothetical phosphorylation sites of MKRP1 and other kinesins, MKRP1 however lacks the C-terminal DUF3490 domain common to HINKEL and NACK1 (Figure 4) and we did not detect HopZ1a acetylation at the C-terminus of HINKEL.

Additional acetylation sites may exist on HINKEL and MKRP1 (and HopZ1a) since LC-MS/MS analysis is unable to detect all peptides generated from trypsin digests of the proteins of interest; we only acquired 47-51% coverage of HINKEL, 56-63% coverage of MKRP1, and 43-55% coverage of HopZ1a (Figure 4). Thus, our acetylation analysis is conservative and it remains possible that HopZ1a acetylates additional residues of HINKEL and/or MKRP1 that we were unable to observe. Although HINKEL is acetylated within its kinesin motor domain at positions S88 and T90, the corresponding residues were not acetylated in MKRP1. The acetylation sites of MKRP1 are not present in HINKEL (not shown) and HINKEL has a C-terminal DUF3490 domain that is absent from MKRP1 (Figure 4). Thus, if acetylation of these two kinesins is an important function of HopZ1a *in planta*, they are likely to be regulated by contrasting mechanisms. Nevertheless, these data indicate that HopZ1a can target *A. thaliana* Kinesin 7 family members.

We had hoped to demonstrate functional importance for HopZ1a acetylation of kinesins by comparing the growth of *P. syringae* in transgenic *Arabidopsis* plants that conditionally-overexpress either wild type HINKEL or putative acetyl-mimetic alleles (with glutamine substitutions at S88 and T90). However, despite independent cloning and transformation attempts by many of us (AHL, DPB, TL, and JZ), we were consistently unable to detect even modest expression of HA-tagged recombinant HINKEL in *Agrobacterium*-transformed *Arabidopsis* (data not shown). Although the dexamethasone-inducible expression vector we employed (Aoyama and Chua 1997) has been used successfully in the past by ourselves and by others, it is known that leaky (uninduced) expression levels from the dexamethasone-dependent promoter are not insignificant (data not shown). Coupled with an absence of available T-DNA knockout lines for *HINKEL* and *MKRP1* from the Arabidopsis Biological Resource Center (https://abrc.osu.edu/) (de Lucas et al. 2014), our present inability to overexpress HINKEL suggests that *Arabidopsis* plants are very sensitive to mis-regulation of these proteins and that endogenous regulatory mechanisms robustly prevent their overexpression. Although this complicates assessment of the *in planta* significance of kinesin acetylation by HopZ1a, overall our efforts highlight the potential offered by integrating functional genomic and physical interaction datasets to aid virulence target inference for T3SEs from important pathogens like *P. syringae*.

## ACKNOWLEDGEMENTS

This work was supported by Natural Sciences and Engineering Research Council of Canada awards to D.S.G. and D.D; a Canada Research Chair in Plant-Microbe Systems Biology (D.D.) or Comparative Genomics (D.S.G.); the Centre for the Analysis of Genome Evolution and Function (D.D. and D.S.G.). This study is done in memory of Dr. Jianfeng Zhang.

